# Positive selection and gene duplications in tumour suppressor genes reveal clues about how cetaceans resist cancer

**DOI:** 10.1101/2020.01.15.908244

**Authors:** Daniela Tejada-Martinez, João Pedro de Magalhães, Juan C. Opazo

## Abstract

Cetaceans are the longest-living species of mammals and the largest in the history of the planet. They have developed mechanisms against diseases such cancer, although the underlying molecular bases of these remain unknown. The goal of this study was to investigate the role of natural selection in the evolution of 1077 tumour suppressor genes (TSGs) in cetaceans. We used a comparative genomic approach to analyse two sources of molecular variation in the form of dN/dS rates and gene copy number variation. We found a signal of positive selection in the ancestor of cetaceans within the *CXCR2* gene, an important regulator of DNA-damage, tumour dissemination, and immune system. Further, in the ancestor of baleen whales, we found six genes exhibiting positive selection relating to such diseases as breast carcinoma, lung neoplasm (*ADAMTS8*) and leukaemia (*ANXA1*). The TSG turnover rate (gene gain and loss) was almost 2.4-fold higher in cetaceans as compared to other mammals, and noticeably even faster in baleen whales. The molecular variants in TSGs found in baleen whales, combined with the faster gene turnover rate, could have favoured the evolution of their particular traits of anti-cancer resistance, gigantism and longevity. Additionally, we report 71 genes with duplications, of which 11 genes are linked to longevity (e.g. *NOTCH3* and *SIK1*) and are important regulators of senescence, cell proliferation and metabolism. Overall, these results provide evolutionary evidence that natural selection in tumour suppressor genes could act on species with large body sizes and extended life span, providing novel insights into the genetic basis of disease resistance.

## Background

Why do not larger organisms always have a higher risk of developing cancer than smaller ones? This is an unresolved question that was first observed by Richard Peto in 1977 [1] which later became known as “Peto’s paradox” [2,3]. This paradox is based on the observation that the risk of developing cancer should increase with the number of cells. The higher number of cells the greater the number of cell divisions and the higher the probability of DNA damage and the transformation of a normal cell into a cancerous one [4,5]. Accordingly, we should expect a higher probability of developing cancer in larger and longer-lived organisms than smaller ones, although this does not occur [6–10]. Several hypotheses have been proposed to explain the different evolutionary strategies that have arisen to fight cancer [11], including specific adaptations to the environment, fitness, DNA damage responses, tissue microenvironments, and cellular senescence [12,13]. Recently, the availability of the genomes of a wide variety of species has prompted the search for genes that could explain the differences arising in cancer incidence between species [14]. In particular, understanding the molecular basis of the anticancer mechanisms in large and long-lived species may shed light on several fundamental questions in medicine.

How organisms reach and maintain larger body sizes and/or achieve increased longevity has puzzled scientists for decades [15,16]. For example, the naked mole rat (*Heterocephalus glaber*) has an average body mass of 35g and a lifespan of 35 years [17]. Long-term studies suggest that their mortality rate [18] and the risk of developing cancer does not seem to increase with age [19]. In this species, special mechanisms like early contact inhibition would be, in part, responsible for their extended lifespan [20,21]. Among vertebrates, other anticancer-ageing mechanisms have been described which could potentially explain the maintenance of large body sizes and/or increased longevity [11]. For example, in the little brown bat (*Myotis lucifugus*) telomere dynamics and changes in the genes associated with growth factors are related to repair mechanisms that prevent DNA damage associated with ageing [22,23]. In the yellowmouth rockfish (*Sebastes reedi*), there are no signs of aging in replicative senescence [24], while in the African elephant (*Loxodonta africana*) it is explained as a variation arising in gene copy number in TP53 and LIF genes [25]. Molecular variation in tumour suppressor genes could work in the manner of a compensatory mechanism, providing additional and novel ways to avoid or prevent cancer [4]. However, the evolutionary history of tumour suppressor genes remains unknown.

Cetaceans evolved from a terrestrial ancestor and re-entered into the oceans around 50 million years ago [26]. Their adaptation to an aquatic lifestyle required many morphological, physiological, and behavioural changes. The sub-order Cetacea comprises two main lineages; namely Odontoceti (or toothed whales) and Mysticeti (or baleen whales) [26]. Baleen whales and toothed whales display a vast variation in terms of body mass and maximum lifespan, ranging from 17 years for the Pygmy sperm whale (*Kogia breviceps*) to 211 years in the bowhead whale (*Balaena mysticetus*); and from 50 kg for Maui’s dolphin (*Cephalorhynchus hectori maui*) to 175 tons for the blue whale (*Balaenoptera musculus*) [17]. Because there are species that live over a hundred years, they are exposed for a much longer period to the appearance of harmful mutations, pathogens, or diseases. Thus, it is expected that cetaceans developed improvements in their immune systems, DNA repair mechanisms and other biological processes which could act to favour greater longevity and body mass [27,28]. Further, recent investigations have found a signal for the positive selection of those genes which could be related to the observed variations in terms of body size [27]. However, although in mammals the body of evidence suggests that tumour suppressor genes play a key role in the evolution of body size and longevity by decreasing the incidence of diseases such as cancer [15,29], it still remains an open question as to which molecular variants in cetaceans are responsible for the lower observed incidence of cancer [30–32].

The goal of this study is to investigate the evolution of tumour suppressor genes in cetaceans by evaluating two forms of molecular variation, namely d_N_/d_S_ and gene copy number. We report a signal of positive selection in those genes which are involved in the control of cancer onset and progression. The turnover rate of tumour suppressor genes was faster in cetaceans relative to other mammals. We report 71 genes exhibiting duplications in one or more species of cetaceans which are known to be involved in multiple human disorders and ageing. This approach highlights how studying the role of natural selection within genes associated with human health could lead to advances in our understanding of the genetic basis of disease resistance.

## Materials and Methods

### Relationship between body mass and longevity in cetaceans

We used the AnAge database to evaluate if there exist differences between the longevity and body mass in cetaceans relative to other mammals. The Caper R-package was used to make a linear regression through the Phylogenetic generalized least squares (PGLS) [33] models. PGLS assumes a multivariate Brownian motion process and consider the phylogenetic relationships among species. The phylogenetic tree was obtained from Uyeda et al. (2017) [34].

### DNA sequences and taxonomic sampling

To study the evolution of tumour suppressor genes (TSGs) in cetaceans we implemented a phylogenetic design including 15 mammalian species. Our taxonomic sampling included five Odontocetes (bottlenose dolphin, *Tursiops truncatus*; orca, *Orcinus orca*; beluga, *Delphinapterus leucas*; Yangtze river dolphin, *Lipotes vexillifer*; the sperm whale, *Physeter catodon*); two Mysticetes (common minke whale, *Balaenoptera acutorostrata*; bowhead whale, *Balaena mysticetus*); five other members of the superorder Laurasiatheria (cow, *Bos taurus*; pig, *Sus scrofa*; dog, *Canis familiaris*; horse, *Equus caballus*; microbat *Myotis lucifugus*); two Euarchontoglires (human, *Homo sapiens*; mouse, *Mus musculus*); and one Atlantogenata (African elephant, *Loxodonta africana*). The coding sequences for each species were downloaded from Ensembl v.96 (http://www.ensembl.org), NCBI database [35] and the Bowhead whale genome project (http://www.bowhead-whale.org/) (supplementary table 1). To remove the low-quality records, sequences were clustered using CD-HITest v.4.6 [36] with a sequence identity threshold of 90% and an alignment coverage control of 80%. After that, the longest open reading frame was kept using TransDecoder LongOrfs and TransDecoder-predicted in TransDecoder v3.0.1 (https://github.com/TransDecoder/TransDecoder/).

**Table 1.** Human diseases related to genes with a signature of positive selection in the ancestor of cetaceans and baleen whales. The information was obtained from GeneCards (https://www.genecards.org/),

| Branch |  | Diseases associated |
| --- | --- | --- |
| **Ancestor Cetacea** | CXCR2 | Autosomal recessive severe congenital neutropenia – Neutrophil migration - Immune system |
| **Ancestor Mysticeti** | ADAMTS8 | Diseases of glycosylation – Lung neoplasm |
|  | ANXA1 | Anti-inflammatory activity. Hairy Cell Leukemia and Penis Squamous Cell Carcinoma. |
|  | DAB2 | Malignant epithelial mesothelioma – Hypercholesterolemia autosomal recessive - ovarian carcinoma |
|  | DSC3 | Hypotrichosis - Recurrent skin vesicles - Subcorneal pustular dermatosis |
|  | EPHA2 | Cataract 6, multiple types - Early-Onset posterior subcapsular cataract |
|  | TMPRSS11A | Iron-Refractory iron deficiency anemia – Ichthyosis follicular |

### Homology inference

We inferred homologous relationships between the 1077 tumour suppressor genes described for humans, which are available from publically accessible databases (Tumor Suppressor Gene Database, https://bioinfo.uth.edu/TSGene/ and the Tumor Associate Gene, http://www.binfo.ncku.edu.tw/TAG/GeneDoc.php) (supplementary table 2), and the other 14 species included in our study using the program OMA standalone v.2.3.1 [37]. OMA standalone is a graph-based method that infers homology using strict pairwise sequence comparisons “all-against-all” based on evolutionary distances. It also has a survey of the relationships between species, which for this study were included *a priori*. We inferred two types of groupings: 1) OMA Groups (OG), containing the sets of orthologous genes, and 2) Hierarchical Orthologous Groups (HOGs). Briefly, tHOGs are those genes that have descended from a single ancestral gene within a species against which gene duplications and losses can be contrasted. This algorithm has previously shown high precision, minimising the errors in orthology assignment in comparison with other homology inference methods [38,39]. Using these HOGs groups, the number of copies of tumour suppressor genes was obtained on a per species basis. We used a home-made script as an extra filter so as not to overestimate the number of copies of TSG. For each HOG we filtered the longest transcript given the same GeneID (supplementary file 5).

The amino acid sequences were aligned using the L-INS-i algorithm from MAFFT v.7 [40]. Nucleotide alignments were generated using the amino acid alignments as a template using the function pxaa2cdn in phyx [41]. Finally, to reduce the chance of false positives arising within low-quality alignment regions, we added a cleaning step using the codon.clean.msa algorithm of the rphast package [42] with the associated human tumour suppressor gene as reference sequence.

### Natural selection analysis

To evaluate the role of natural selection in the evolution of tumour suppressor genes, we employed the codon-based model within a maximum likelihood framework using the program PAML v4.9 [43], as it is implemented in the ETE-toolkit with the ete-evol function [44]. We used the branch-site model to estimate the signature of positive selection with the last common ancestor of Cetacea, Mysticeti (baleen whales), and Odontoceti (toothed whales). We compared the null model, wherein the value of ω in the foreground branch was set to 1, with the model in which the omega value was estimated from the data using the likelihood ratio tests (LRT) [45]. We applied the false discovery rate (FDR) [46] as a multiple tests correction. The genes with an FDR<1 were categorised with a positive selection. Finally, we used branch models to compare the dN/dS distribution of TSGs between cetaceans and other mammals. We implemented a model in which an omega value was estimated for cetaceans and another for all other branches of the tree. To estimate whether the omega distributions were significantly different, we used the Kolmogorov-Smirnov test in R (www.r-project.org/).

### Gene copy number variation analysis

To estimate the gene turnover rate of the tumour suppressor genes we used the software CAFE, v. 4.2.1 (Computational Analysis of Gene Family Evolution) [47]. CAFE uses a stochastic birth and death model to estimate the expansions and contractions of genes using, as a frame of reference, sister group relationships arising among species relative to divergence times. Using this approach we can infer a maximum likelihood value for the rate of evolution (λ) and the direction of the change in terms of the size of the gene families across different lineages with a *p*<0.01. The *p*-value describes the likelihood of the observed number of genes appearing given average rates of gain and loss. All the comparisons were calculated with a genome assembly and annotation random error, assuming that all branches in the tree share the same unique λ. The error model was calculated using the caferror.py script in CAFE [47]. The estimated error value = 0.06875 was included in the further estimations. The family-wide *p*-value describes the likelihood of the observed number of genes arising, given average rates of gain and loss (please see supplementary files 3-4). We implemented two independent models. In the first, one λ was estimated for cetaceans as a total group and the other for the outgroup. In the second model, we estimated four λ values for (i) the stem cetácea; (ii) the total group of Mysticeti; (iii) the total group of Odontoceti; and (iv) the outgroup, respectively. The divergence time between species was obtained from the TimeTree database (http://www.timetree.org/) [48]. However, it should be noted that Odontoceti exhibit a polytomy, so we then re-calibrated the tree using the R packages APE and Phytools [49,50] with the estimated ages and a time-calibrated phylogeny for Odontoceti [51].

### Enrichment analysis

The enrichment analysis was performed using the R package WebGestaltR v0.4.4 [52]. The set of duplicated genes were tested for significant pathway associations using the hypergeometric test for Over-Representation Analysis (ORA) [53]. We selected the categories of gene ontology biological processes, diseases OMIM, and human phenotype and we considered overrepresented categories to be those with a significance level above that of an FDR of 0.01 after correction with the Benjamini-Hochberg multiple test. The enrichment analysis was made independently using as a background the human protein-coding genes and relative to the 1,077 TSGs. Finally, we overlapped the TSGs in terms of positive selection and copy number variation with the GenAge database (build 19,307 genes) [17] and CellAge Database of Cell Senescence Genes [54].

To test if the TSGs with duplications are involved in the longevity process more than would be expected by chance, we performed a gene overlap analysis using the GeneOverlap package in R [55]. The strength of the association was measured via the odds ratio, wherein an odds ratio < 1 implies no association between the lists; and an odds ratio > 1 infers a strong association. Protein coding genes from humans were used as a background (20,183 genes). To make the comparison, we merged the list of genes from the GenAge database and the CellAge Database of Cell Senescence Genes as the “longevity associated genes” and we also employed the list of 1,045 TSGs obtained from the Hierarchical Ortho Groups (HOG) as well as the list of 72 TSGs.

## Results

### Relationship arising between body mass and longevity in cetaceans

We compared the relationship between longevity and body mass in both cetaceans and other mammals. The PGLS analysis (see Materials and Methods), using data obtained from the AnAge database, reveals that the relationship is significantly different across cetaceans in comparison to other mammalian species (*p*-value: < 2.2e-16, Residual standard error: 0.04185 on 929 degrees of freedom, multiple R-squared: 0.08159, adjusted R-squared: 0.07862, figure 1).

**Figure 1.**
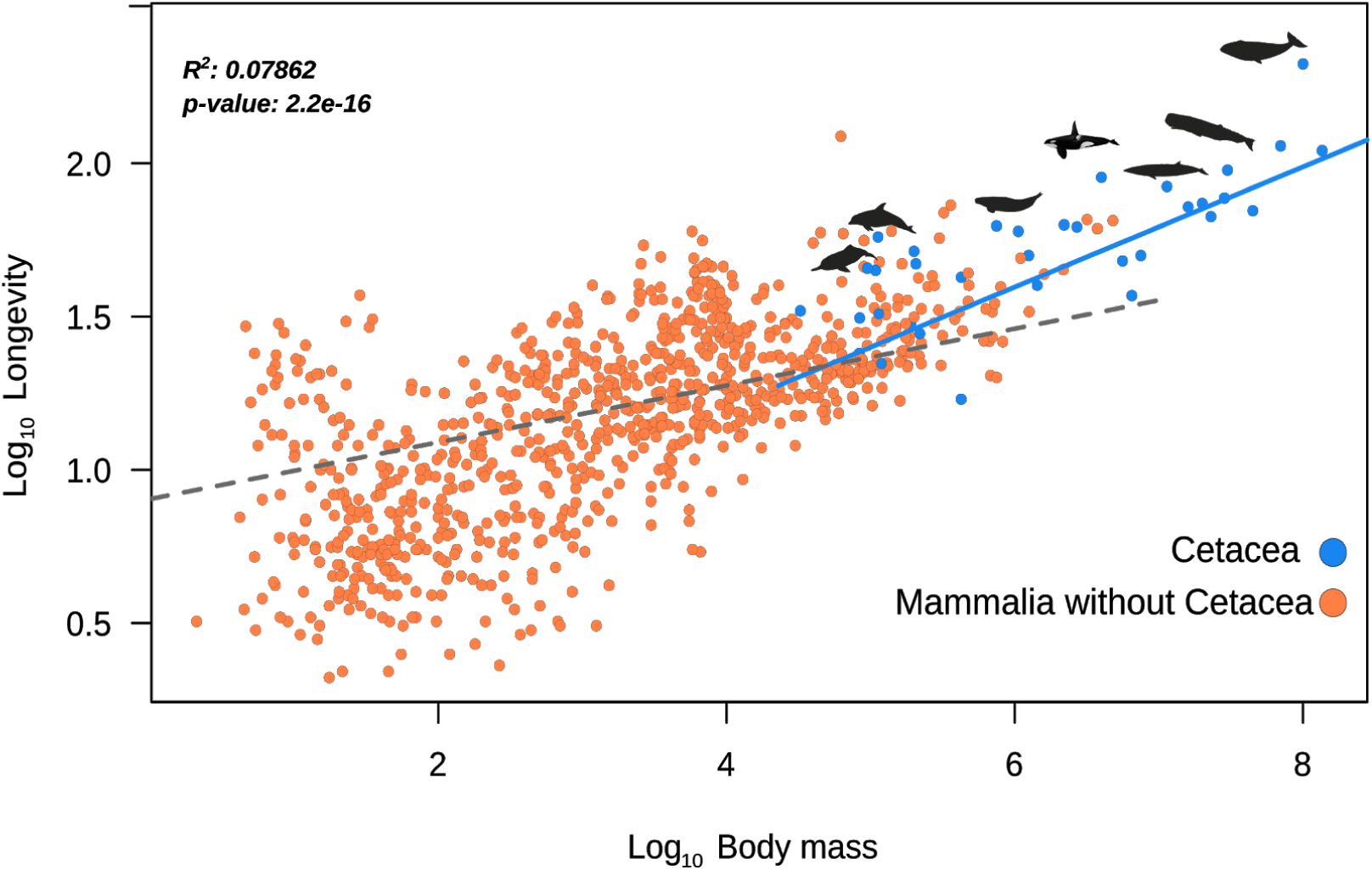
Phylogenetic generalized least squares (PGLS) between the body mass and longevity across 930 mammalian species. The results from the regression showed that the relationship between body mass and longevity is significantly different in cetaceans as compared to other mammals (*p<*0.001).

### Homology inference

We performed an evolutionary analysis to understand the role of natural selection in the evolution of tumour suppressor genes in cetaceans. Employing the 1,077 TSGs described for humans, we obtained 362 OrthoGroups (OG) from the 15 mammalian species included in this study (supplementary file 1) as well as 1,044 hierarchical orthologous groups (HOG) containing two or more species (supplementary file 2).

### TSGs with positive selection in cetaceans are correlated with multiple human disorders

Within the 362 orthologous genes selected, we found that a signal for positive selection within 7 tumour suppressor genes (figure 2a, supplementary table 3). We report that the *CXCR2* gene exhibits a signature of positive selection in the ancestor of cetaceans and also in the genes *DAB2, ADAMTS8, DSC3, EPHA2, TMPRSS11A, ANXA1* in the ancestor of baleen whales. The GeneCards - Diseases associated database showed that those genes with the signature of positive selection are implicated in multiple forms of cancer, such as lung neoplasm (*ADAMTS8*), leukemia (*ANXA1*), and teratocarcinoma (*DAB2*) as well as in immune system disorders such as congenital neutropenia syndrome (*CXCR2*) (table 1). In the last common ancestor of baleen whales, a lineage that includes the largest and longest-lived species of mammals, the most represented functional categories were all related to cell cycle regulation, metabolism and apoptosis. It is noteworthy to mention that several genes with the signature of positive selection are related to abnormalities in scalp hair and blood vessel development, phenotypes associated with the evolution of aquatic life. For the branch model, we found that the distribution of the omega values arising between groups is significantly different (D = 0.32764, *p*-value < 2.2e-16), while the kernel-smoothed density estimates of the omega values show that crown cetacea have a distribution of the mean that is greater than for the outgroup (figure 2b - supplementary table 4).

**Figure 2.**
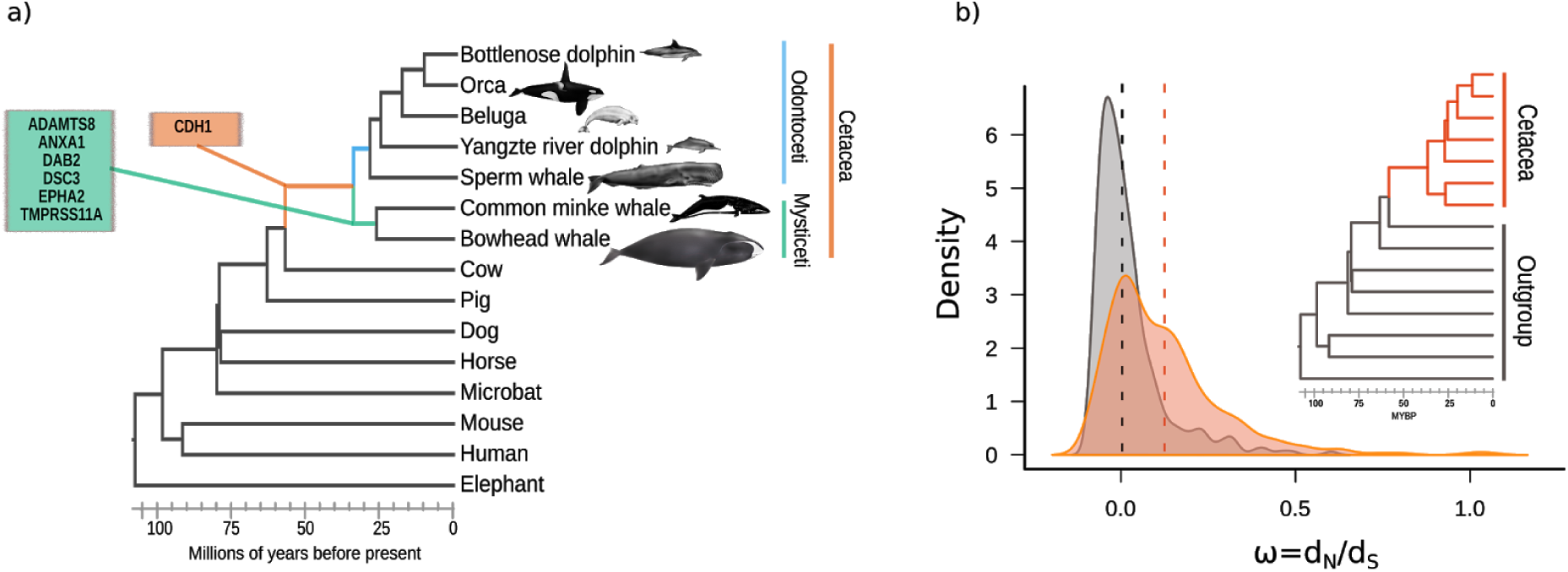
Codon-based models of evolution (ω=dN/dS models). a) Tumour suppressor genes with the signature of positive selection in different branches of the cetacean tree of life. b) Distribution of the omega values estimated separately for cetaceans and all other branches of the tree. The height of the curves is relative to the number of genes with the observed omega values and the dotted lines represent the mean value for cetaceans (orange) and the outgroup (grey). The Kolmogorov-Smirnov test showed that the distributions have statistically significant differences (*p*-value < 0.01).

In summary, we report signatures of positive selection arising in genes that are involved in multiple types of cancer and also in immune system disorders that could increase predisposition to cancer. Since we found more TSGs with the signature of positive selection in the ancestor of baleen whales, we suggest that these lineages could have evolved with a selection for additional anticancer/ageing mechanisms, thereby reducing the risk of cancer and thus favouring longevity and greater body mass. Studying ageing with such a comparative perspective offers an avenue that could potentially provide information about the role of natural selection in species which have found a way to avoid diseases commonly associated with ageing, such as cancer.

### Cetaceans have an accelerated gene turnover rate in comparison to other mammals

The copy number variation analysis revealed that in the first model, the turnover rate of TSGs of cetaceans, as a total group, is more than two times faster (λc = 0.00074) than the rest of the tree (λo = 0.00031) (figure 3a). In the second model, in which we specified four rates of gene family evolution, we found that the turnover rate values estimated for the last common ancestor of baleen whales (λ_my_= 0.0019) was faster than for the last common ancestor of cetaceans (λ_C_= 0.00022) or the last common ancestor of toothed whales (λ_od_= 0.00015) (figure 3b).

**Figure 3.**
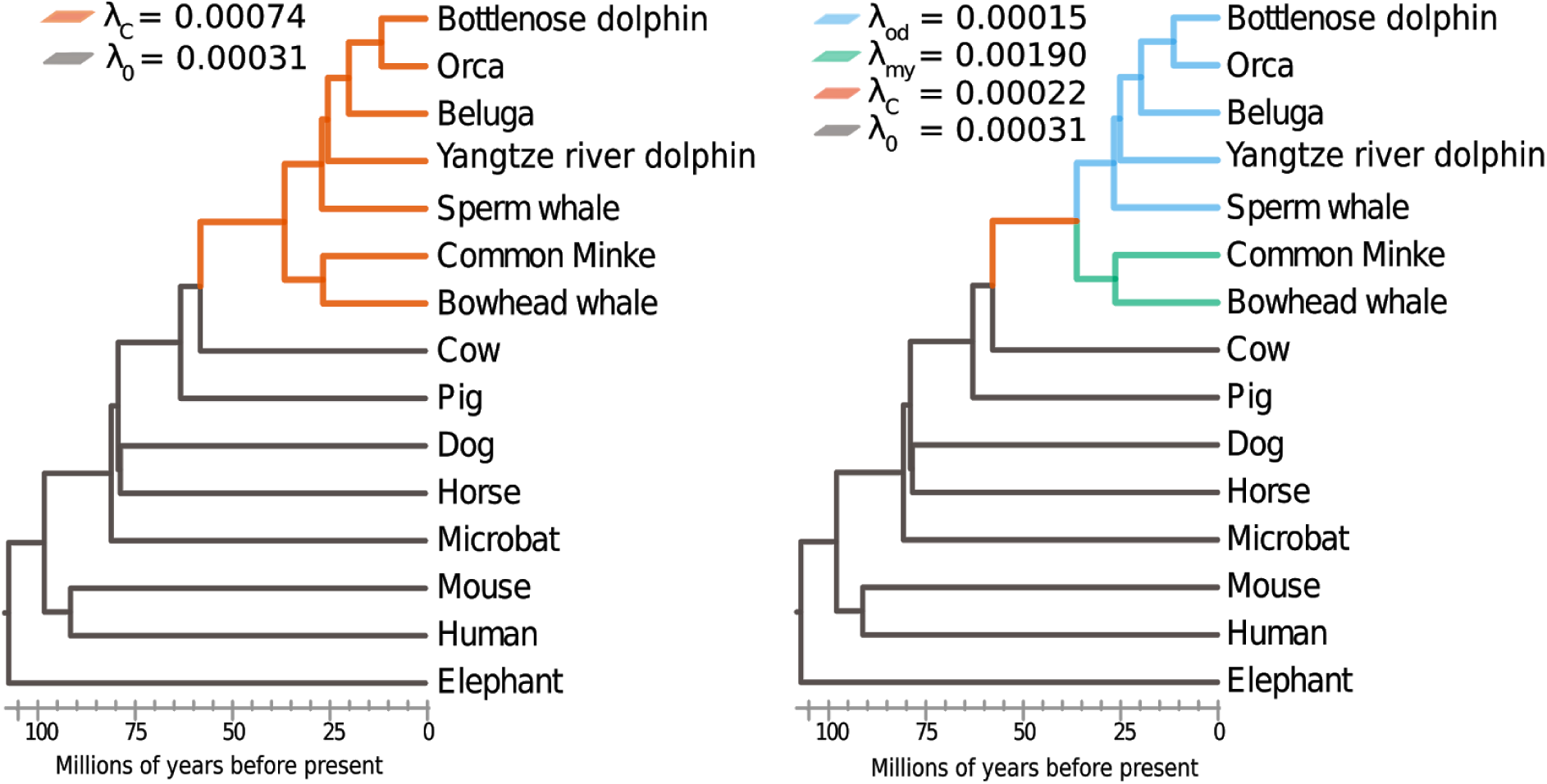
Turnover rates of tumour suppressor genes (TSGs) in cetaceans. a) The first model represents the rate of evolution (λ) of TSGs for the total group of cetacea (orange) and the rate of evolution of TSGs for the outgroup (grey). The *λ* values show that the gene turnover rate of TSGs in cetaceans is almost 2.4 times faster than for other placental mammals. b) In the second model, the λ values represent the rate of evolution for the total group of baleen whales (green), the total group of toothed whales (light blue), and for the ancestor of all cetaceans (orange). The gene turnover rate in the total group of baleen whales is faster in relation with other groups.

To gain further insights into those mechanisms pertaining to ageing and disease resistance which are associated with the tumour suppressor gene families, we identified those TSGs with specific duplications in cetaceans. According to our analyses, we identified that 71 TSGs were duplicated in one or more cetacean species (figure 4a; supplementary table 5-6). Among those genes with duplications, we found statistical significance for the involvement of 11 TSGs in longevity processes (figure 4c; supplementary table 7-8); namely, EEF1A1, H2AFX, HSPD1, MAPK9, GSTP1, PTPN11 (which are known to be involved in ageing); and IGFBP3, NOTCH3, IGFBP3 and SIK1 (known promoters of senescence); while RBX1 and YAP1 are known to inhibit senescence. The results from the enrichment analysis, using the 1,077 TSGs as a reference gene list, infer no significant matches below threshold (fdr 0.1), meaning that the biological functions in which those genes with duplications are involved are not significantly overrepresented relative to other TSGs. However, the enrichment analysis made using the human protein-coding genes revealed that they are significantly enriched within several biological processes (figure 4b, supplementary table 9) such as cell proliferation, cell migration, the cellular apoptosis response to epidermal growth factor stimulus, and metabolic processes. We also report that 17 genes which are duplicated are involved in neurogenesis (e.g. *MAPK9, FAT, EFNA5, NOTCH3* and *YAP1*) or neurological disorders such as the Arnold-Chiari malformation (*DKK1, MAF, NOTCH3, SETD2*).

**Figure 4.**
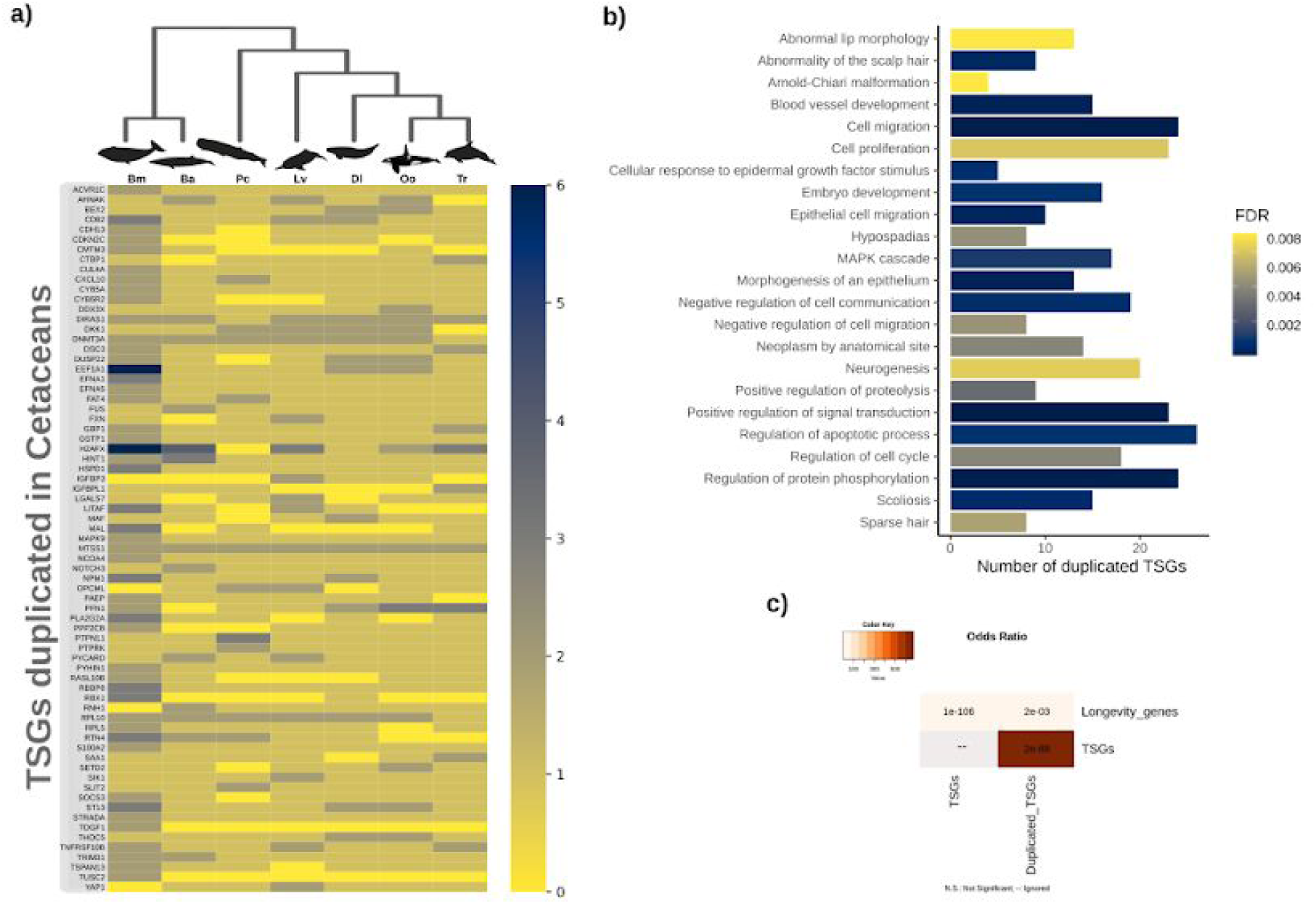
Tumour suppressor genes (TSGs) with duplications in cetaceans. a) Heat map showing copy number variation of TSGs in cetaceans. The colour code corresponds to the number of copies. Key: Tt -*Tursiops truncatus*, Oc-*Orcinus Orca*, Dl-*Delphinapterus leucas*, Lv-*Lipotes vexillifer*, Pc-*Physeter catodon (*synonym name of *Physeter macrocephalus)*, Ba-*Balaenoptera acutorostrata* and Bm-*Balaena mysticetus. b)* Enrichment analysis for the duplicated genes found in cetaceans using the whole genome as background. The image shows the first 30 categories with an FDR<0.01 for biological processes, Diseases OMIM, and Human phenotype ontology. c) Overlap analysis showing that those TSGs with duplications are involved in the longevity process more than would be expected by chance (*adjusted p-value* <0.01).

In summary, we uncovered an accelerated rate of evolution of TSGs among cetaceans in comparison to other mammalian species. Further, we report 71 gene families with duplications relating to other important anticancer processes, such as DNA repair, metabolism, apoptosis, ageing and senescence. Intriguingly, these have also been reported in other long-lived species.

## Discussion

Cetaceans are the longest-lived mammals and cancer-resistant species. Our comparative analysis revealed that the lineage of cetaceans generally lives longer relative to body mass in comparison to other mammals. We report seven TSGs exhibiting positive selection, one within the ancestor of the group and six within the ancestor of baleen whales, suggesting that baleen whales could have evolved additional anticancer mechanisms. Moreover, the turnover rate of gene gain and loss (*λ*) was accelerated in the cetaceans relative to other mammalian species and we report that 71 TSGs were duplicated in one or more cetacean species. To further understand the evolution of TSGs, we studied the biological processes in which those TSGs exhibiting positive selection and duplications are involved, with a specific focus on important anti-cancer, longevity and senescence pathways. In the ancestor of all cetaceans, we report the signature of positive selection in the *CXCR2* gene, known to regulate the recruitment of leukocytes during inflammatory processes. This gene has also been described as a crucial factor in tumour cell dissemination [56,57], immune system disorders such as congenital neutropenia [58], and senescence [59]. In those species that evolved large body masses, replicative senescence is considered to be an important tumour suppressor mechanism [21].

The fact that baleen whales have more positively selected TSGs and a faster gene turnover rate could be interpreted as translating into stronger protective anticancer and ageing mechanisms which might have favoured the evolution of gigantism and longevity amongst cetaceans. From those genes exhibiting positive selection in baleen whales, the genes *DAB2* and *ADAMTS8* have previously been implicated in breast cancer [60,61]; while *ADAMTS8* is also involved in angiogenesis and linked to several other forms of cancer, including brain tumours [62], also serving as a promising biomarker in lung neoplasia [63]. On another hand, *ANXA1* is an important anti-inflammatory gene related to leukemia and is known to be involved in cellular proliferation, differentiation and apoptosis [64]. Considering the evolution of gigantism in baleen whales, molecular variants involved in angiogenesis, inflammatory processes, and apoptosis may have arisen as a protective mechanism against the development of tumours.

Recent studies of the genome of the humpback whale (*Megaptera novaeangliae*) also suggested that the evolution of body mass in cetaceans could be related to strong selection in terms of cancer resistance pathways and large segmental duplications [28]. Here we discovered several TSGs exhibiting duplication in cetaceans, an observation that could help future investigations to solve the secrets underlying their longevity and cancer resistance. For example, we found two copies of the *MTSS1* gene, implicated in cell proliferation, not only within all cetaceans but also in other several species from the outgroup. Other important genes identified were *RPL10*, which is involved in the regulation of apoptosis and metabolic processes, and *DNMT3A*, implicated in neurogenesis and scoliosis, with duplications present within all cetaceans species except for the common dolphin. In fact, in this study, the species with the most gene duplications is the bowhead whale, the only mammal that is recognised as living for more than two centuries. In contrast, the bottlenose dolphin, with fewer duplicated TSGs has the highest incidence of cancer identified within cetaceans [65,66]. However, the possibility remains that the differences arising between cancer incidence and longevity within cetaceans may arise due to a reduction in the number of duplicated TSGs (as in the bottlenose dolphin). Moreover, it is also likely that there is a sampling gap which indicates that more studies are required. For example, in the bowhead whale genome, further genes were identified with the signature of positive selection that are known to be involved in DNA repair, cell cycle regulation, resistance to ageing, and cancer [67]. It has been suggested that increasing the number of copies could serve a protective ‘guardian’ effect in preventing somatic mutations from spreading through the cellular population [6,68,69]. It would be interesting to investigate the effect of those conserved TSGs variants identified in whales in human cells. Such an avenue of investigation could eventually lead to an understanding of novel pathways that serve to improve the cellular mechanisms protecting against DNA damage, cancer and ageing.

Ageing is the single largest risk factor for the development of cancer and is characterised by the progressive accumulation of cellular damage, yet the mechanisms linking ageing and cancer remain unclear [70]. We report that the turnover rate of TSGs in cetaceans is almost 2.4-fold faster than for other mammals. This acceleration in turnover rate is associated with several TSG duplications, among which eleven genes are known to be associated with longevity and the ageing process. Previous studies have shown that longevity-associated gene families evolve faster (as measured by an increase in the number of gene family members) in long-lived species [71]. One of the main processes which has already been demonstrated to play an important role in ageing and cancer suppression is the irreversible arrest of normally dividing cells, or cell senescence [72]. From those duplicated genes in cetaceans, *IGFBP3, SIK1* and *NOTCH3* are known senescence promoters. *IGFBP3* is a bone health determinant in middle-aged and elderly European men [73] and plays an important in cell proliferation, apoptosis [74], and breast cancer [75]; while *SIK1* expression has been correlated with arterial blood pressure, one of the primary pathologies associated with ageing in humans. *SIK1* has also been implicated in sympathetic nervous system hyperactivity in mammals [76]. Similarly, the *NOTCH3* gene, which has two copies in the Minke whale, also promotes cell senescence and has been correlated with the development of brain lesions with the age [77]. The increase in the number of copies of ageing genes in cetaceans could provide additional protection against the development of several types of cancer as well as ageing-related diseases.

Although in this study we primarily focus on the roles of genes in cancer and ageing, it is important to highlight that they could also perform functions related to other phenotypic outcomes such as those associated with phenotypic remodelling in response to the conquest of the aquatic environment. For example, the *DSC3* gene, positively selected in Mysticeti, is a biomarker for colorectal [78] and breast cancer [79]. However, mutations in this gene are also correlated with hair loss disorders [80] such as hypotrichosis [81]. Hair loss in cetaceans was related to their hydrodynamic adaptation in an aquatic environment [82]. In the same vein, several TSGs which have been duplicated in whales (e.g. *ACVR1C, CTBP1, DDX3X*) are involved in metabolic processes that may have responded to those selective pressures involved in energy consumption, diet, and increases in body mass. Furthermore, *ANXA1*, an anti-inflammatory gene, and *EPHA2*, an antibody receptor target in chemotherapy [83], are also involved in blood vessel development, another important feature in underwater evolution [84]. Considering the pleiotropic effects of TSGs, a more comprehensive understanding of the consequences of natural selection at a molecular level in the evolutionary history of cetaceans is required.

## Conclusions

The main goal of this study was to shed light on the evolution of TSGs in cetaceans. We report signals of positive selection in 7 TSGs and an accelerated gene turnover rate in comparison to other mammals, with duplications arising in 71 genes in one or more cetacean species. Those genes with the characteristic signature of positive selection are implicated in multiple diseases, including bladder cancer, breast cancer, leukaemia and immune system disorders (e.g. congenital neutropenia). The turnover rate of TSGs was almost 2.4 times higher in cetaceans as compared to other mammals. From those genes with duplications, 11 are linked to longevity and senescence (e.g. *NOTCH3, IGFBP3*, and *SIK1*). Although the enrichment analysis using the 1077 TSGs as a background did not lead to significantly enriched biological categories, the same analysis using all protein coding genes as a background gives us a sense of which functions are associated with the duplicated genes. Some of those functions included cell proliferation, apoptosis, and metabolism. Remarkably, most of the genes exhibiting evidence of positive selection and duplication were found amongst the baleen whales. Considering that the clade of baleen whales includes the biggest extant species known and that such species are also among the longest living of all mammals, we suggest that they could have evolved with additional mechanisms that conferred both resistance to cancer and longevity. Nonetheless, due to possible pleiotropic effects of TSGs, the genes we identified as duplicated and under positive selection could also have been important in the cetacean evolution to aquatic life. Studying genes that are related to human malignancies from a comparative perspective offers a novel avenue that could provide additional clues about the role of natural selection in the evolution of all species and, potentially, lead to the discovery of novel therapeutic approaches. This study provides evolutionary evidence that natural selection among TSGs could act favourably for those species with larger body sizes and greatly extended life spans, thereby providing insights into the genetic basis associated with the evolution of disease resistance. We also highlight the importance of cetaceans as an evolutionary model to understand the genesis of ageing disorders in humans and other animals.

## Data accessibility

All data generated or analysed during this study are included in this published article and in the supplementary information. The supplementary files 1 and 2 are deposited in the Dryad repository.

## Authors’ contributions

D.T.M and J.C.O conceived the project; D.T.M performed the bioinformatic analysis. D.T.M wrote the manuscript; J.C.O and J.P.M edited the manuscript and were the project advisors.

## Acknowledgements

Members of the Integrative Genomics of Ageing Group at the University of Liverpool, for all the support and advice during DTM’s internship. To Richard Gregory in the Centre for Genomic Research (CGR) and Ian C. Smith in the Advanced Research Computing at the University of Liverpool for their help with computing resources. To Dr. Andres Parada, for the academic discussions and with programming advice. To Dimar Gonzales for the programming advice. To Alejandra Tejada-Martinez for the graphic designs. To Dr. Priyanka Raina for the grammatical revision of the manuscript. Finally to Jesus Tejada and Gloria Martinez for their unconditional support.

